# Registration for Image-Based Transcriptomics: Parametric Signal Features and Multivariate Information Measures

**DOI:** 10.1101/521898

**Authors:** Rebecca Chen, Abhinav B. Das, Lav R. Varshney

## Abstract

Image-based transcriptomics involves determining spatial patterns in gene expression across cells and tissues. Image registration is a necessary component of data analysis pipelines that study gene expression levels across different cells and intracellular structures. We consider images from multiplexed single molecule fluorescent *in situ* hybridization (smFISH) and multiplexed *in situ* sequencing (ISS) datasets from the Human Cell Atlas project and demonstrate a novel approach to groupwise image registration using a parametric representation of images based on finite rate of innovation sampling, together with practical optimization of empirical multivariate information measures.

## I. Introduction

The transcriptome of a cell (or an organism) is the portion of DNA that is expressed as RNA in that cell (or organism). The subset of genes that are expressed in a cell varies depending on cell type and cell state, and recognizing patterns in the transcriptome is an important part of understanding cell function and pathology. The RNA molecules present in a cell can be tagged using fluorescent markers, which can be observed *in situ*, and spatial patterns of gene expression within cells and tissues can be studied [1]. Different combinations of colored fluorescent markers serve as tags for different RNA sequences, and images of different color spectra (multispectral images) must be aligned for analysis.

Researchers have developed methods and algorithms to extract transcript molecule feature sets, localization, and patterns, in tens of thousands of single cells across the human transcriptome [2], [3]. Starfish, a community of computational biologists and software engineers, has developed standardized file formats for input data and analysis [4]. They are building a standard library that consolidates different methods from different steps of the analysis pipeline. The goal is to contribute to the Human Cell Atlas, a project to create comprehensive reference maps of all human cells to understand human health and to diagnose, monitor, and treat disease [5].

An important part of the image-based transcriptomics analysis pipeline is image registration. Image registration in the Starfish pipeline currently addresses pixel-level translational error using an FFT-based phase correlation approach [6]. Phase correlation has also been extended to rotational and scaling error [7] as well as to sub-pixel registration [8]. However, phase correlation does not account for non-linear intensity changes, which can arise when the images to be registered are taken under different conditions (e.g. multimodal, multispectral, different lighting, etc.). To address such intensity nonlinearities, mutual information has been proposed elsewhere as an image similarity measure [9], [10], and under the assumption that images are statistically dependent, we have shown that maximizing mutual information is a theoretically optimal method for registering image pairs [11].

What about registering not just a pair of images but a larger set of images, as in transcriptomics? A natural extension to mutual information is multiinformation, which can be used to jointly register multiple images and has been shown to be superior to sequential pairwise registrations and asymptotically optimal [11]. However, computing the multiinformation of several images is computationally expensive—*O*(*N*^*n*^) for *n* images with *N* pixels. To address this computational issue we develop a novel parametric signal representation for image-based transcriptomics data using finite rate of innovation sampling [12]. We propose a feature- and information-based method which is *O*(*nN*) for *n* images. This algorithm registers images in a joint rather than pairwise manner and has the same output as multiinformation in certain settings when features are properly extracted, see Thm. 1 for a formal statement.

## II. Registration Using Information

### A. Mutual information

Maximizing empirical mutual information for image registration [9], [10] has been commonly used in fields such as medical imaging and remote sensing. Mutual information between random variables *X* and *Y* is defined as

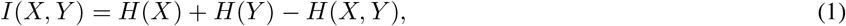

where *H*(*X*) and *H*(*Y*) are marginal entropies and *H*(*X*,*Y*) is the joint entropy of *X* and *Y*. Mutual information can also be formulated as the Kullback-Liebler divergence (relative entropy) between the joint distribution and the product of the marginal distributions:

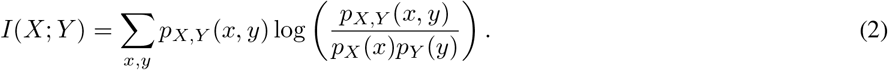

Given two *r*-dimensional images *X*_1_ and *X*_2_ defined over a discrete spatial region Ω, we define an image transformation *T* : Ω → Ω. If *X*_2_ is a transformed version of *X*_1_, we can register *X*_1_ and *X*_2_ by finding

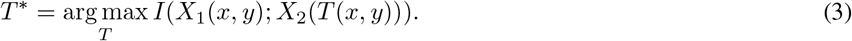

There are several ways to estimate the joint and marginal distributions of images. Two popular ones are the joint histogram method [13] and the Parzen windowing method [9], [14]. After estimating the distributions, maximization problem (3) can then solved using an appropriate optimization algorithm. If properly initialized, a local optimization algorithm such as gradient descent can be used [13]. Global optimization algorithms such as genetic algorithms, simulated annealing, and particle swarm optimization have also been used to maximize mutual information [15]–[18].

### B. Multiinformation

Several groups have proposed using multiinformation to perform groupwise registration [19]–[21]. Guyader et al. show that groupwise multiinformation yields better registration performance than pairwise mutual information for medical imaging settings. Multiinformation, like mutual information, is defined as the KL divergence between a joint distribution and a product of marginal distributions. Given *n* random variables *X*_1_, *X*_2_, …, *X*_*n*_, multiinformation is defined as

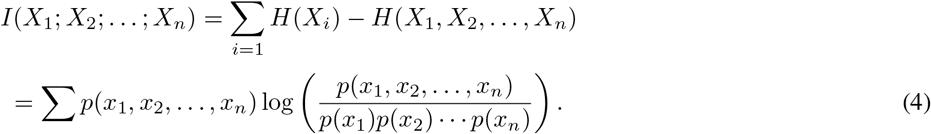

To register *n* images *X*_1_, *X*_2_,…, *X*_*n*_, we find the transformations that maximize multiinformation:

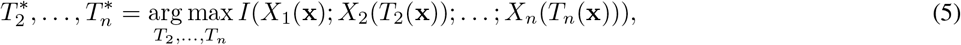

where x = (*x*, *y*) for a 2-D image. Thus a single optimization can be used to register any number of images. We call this the *MM algorithm*. We have recently shown that MM is exponentially consistent for image registration (probability of error goes to 0 exponentially fast as number of pixels goes to infinity) [11]. In fact, we showed that MM is asymptotically optimal in the sense of achieving the same error exponent as maximum likelihood registration. Thus

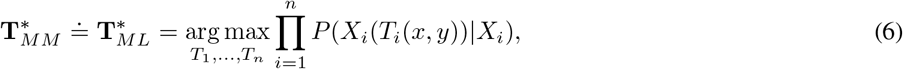

where “≐” indicates equivalent error exponents.

Guyader et al. assume images are jointly Gaussian, which greatly simplifies the empirical distribution estimation [19]. This is usually not a valid assumption for natural images or image-based transcriptomics (although they show that it works for the images they tested). Kern et al. [20] use the approximation

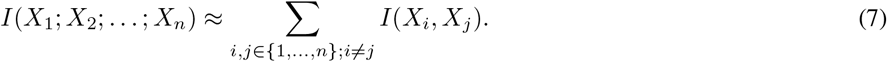

This approximation can greatly overestimate multiinformation in cases where mutual information of image pairs is large.

Rather than approximating the multiinformation function, we initially implement the original objective in (5) using full image histograms. We perform our optimization using particle swarm optimization [22]. The details of our implementation are covered in Sec. IV.

## III. Feature-Based Registration

### A. Background and related work

While MM performs groupwise registration optimally [11], it is computationally expensive. Feature-based algorithms tend to be more efficient. Many registration methods first extract features and then match only the extracted features across images. Others have used corner detectors and edge detectors for registration [23], [24]. Phase correlation in the Fourier domain is applied to these features to retrieve translations and rotations. A particularly successful method is based on Lowe’s scale-invariant feature transform (SIFT) [25]. SIFT extracts local scale-invariant features, called keypoints, from images. Key points are image features that are visually interesting (e.g., corners and curved edges), and feature matching and clustering is performed to detect and register objects. Numerous variants of SIFT have been used in multi-modal image registration (we reference just a few) [26]–[29].

Baboulaz and Dragotti formulate feature-based registration as a finite rate of innovation problem, with applications in super-resolution [30]. They use a Canny-like edge detector and use the intersections of those edges (corners) as features. They then match corners across images using correlation and RANSAC methods (also used in SIFT) and demonstrate that exact registration is possible using only the detected corners.

We apply finite rate of innovation sampling to multispectral registration by representing smFISH images as sums of delta functions. We demonstrate that these features alone are enough to perform registration. We also use this sparse representation in conjunction with MM to perform groupwise registration.

### B. Finite rate of innovation sampling

Certain signals that have a *finite rate of innovation* (FRI) [12] can be written in the form

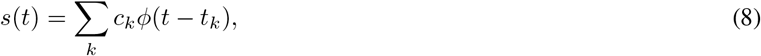

where *ϕ*(*t*) is a known kernel and the number of *t*_*k*_ values per unit time (the rate of innovation) is finite. Then the *innovative* part of the signal lies in *c*_*k*_. Given *c*_*k*_ and *t*_*k*_, we can reconstruct *s*(*t*). Various kinds of filters can be used to identify the innovative part of the signal, and the signal can be perfectly reconstructed using just these samples [31], [32]. Examples of FRI signals include delta trains and piecewise polynomials.

For a 2-dimensional image, we can write (8) as

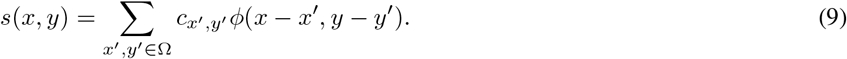

Baboulaz and Dragotti register multiview images using FRI sampling. They model images as sums of polynomials, using a B-spline sampling kernel [30]. The images captured by digital imaging technologies result from the point spread function (PSF) of a lens, which can be used as the sampling kernel *ϕ*(*x*,*y*) in (9). An object in space *o*(*x*,*y*) is filtered through the lens as

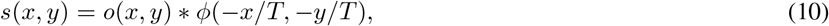

where *T* is the sampling period. For registration, we must obtain the features of *o*(*x*,*y*) from *s*(*x*,*y*). This can be done by deconvolving *s*(*x*,*y*) with the PSF of the imaging system to give *ô*(*p*, *q*). The image *ô*(*x*,*y*) is then processed to extract the features of interest, giving us a weighted sum of spikes:

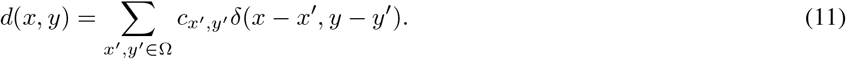

For every feature, there will be a corresponding spike in each misaligned image. To perform groupwise registration, we group these features across images and find the transformations so that spikes corresponding to a feature have the same location in each image.

## IV. Methods and Experiments

In this section, we demonstrate our algorithmic approach on manually shifted and rotated multispectral images, and then on a single smFISH image which we manually shift and rotate. [^1^ Code can be found at https://github.com/toby2476/fishregister] Fig. 2 shows some example smFISH images. Columns correspond to a color channel (there are three total), while rows corresponds to a round of imaging. Disturbances between imaging rounds may cause images to shift, and image registration is necessary. We perform a maximum intensity projection across channels to obtain one image per imaging round, and we perform registration with these images. In Sec. IV-C we present our results on misaligned smFISH images, for which we do not have ground truth data.

**Fig. 1.**
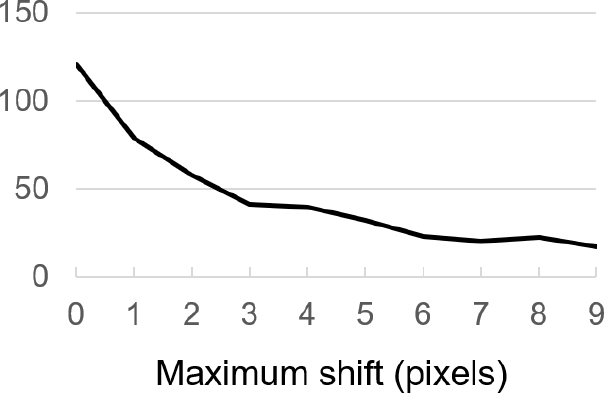
Multiinformation of 30 multispectral images as a function of random shifts.

**Fig. 2.**
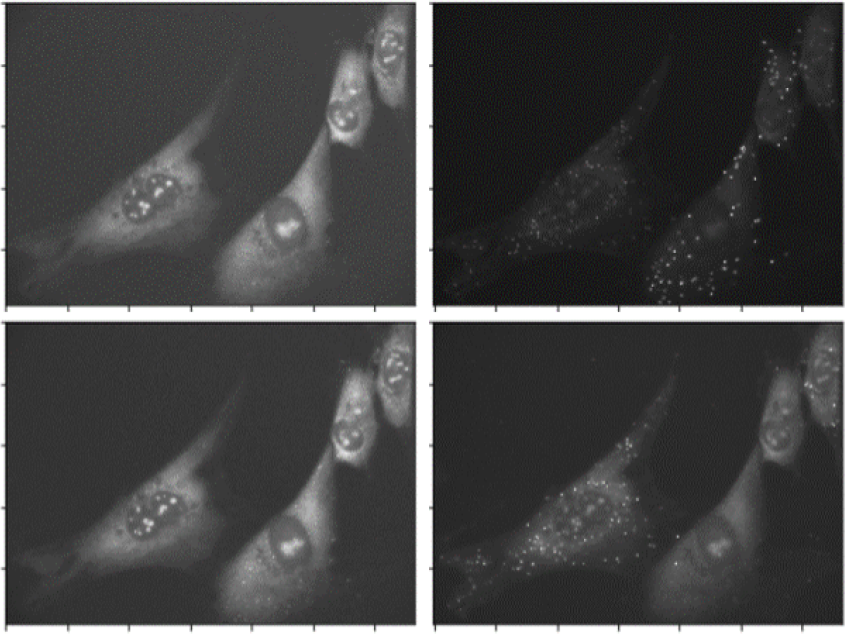
Example smFISH images. Columns are different color channels. Rows are imaging rounds.

### A. Maximizing multiinformation

We implement MM using particle swarm optimization (PSO). To maximize (5), we must search over the space *T*_2_ × *T*_3_ × ··· × *T*_*n*_, where each *T*_*i*_ can have multiple degrees of freedom. For example, if we restrict *T* to include only vertical and horizontal translations, each image has two degrees of freedom, giving us a 2^*n*−1^-dimensional search space. If we include rotations, it becomes a 3^*n*−1^-dimensional search space. We can also include scaling, shearing, etc. Clearly (5) is much more difficult to optimize than (3). We plot the multiinformation of 30 multispectral images from the CAVE database [33] with random horizontal and vertical shifts, estimating distributions using histograms (see Fig. 1). The maximum shift *M* is given on the *x*-axis, and images were shifted with a *x*-shift and a *y*-shift drawn uniformly on [0, *M*]. This matches the plots in [20] showing that mutual information of two images is maximized when they are properly aligned.

To test registration, we use five multispectral images, shown in Fig. 3. We randomly rotate them with angles between [−5, 5] degrees. We find the empirical image intensity distributions for each image using image histograms, and compute the empirical joint distribution using a joint histogram. Using numerous histogram bins causes joint probabilities to become too small, so we use four bins. We calculate entropies and joint entropies using the empirical distributions, obtain multiinformation using (2), and maximize (3) using PSO. Because histograms are computationally expensive, we resized the images to decrease image sizes. We begin with exhaustive search of multiinformation over all rotations and apply the transformation that maximizes multiinformation. (We restrict to rotations—only one degree of freedom per image—so we can test exhaustive search.) Comparing this to PSO yields the same results. Fig. 4 shows images transformed with shifts drawn uniformly from [−10,10] pixels and rotations drawn uniformly from [−10,10] degrees. This example was initialized with 300 particles and converged after 211 iterations.

**Fig. 3.**
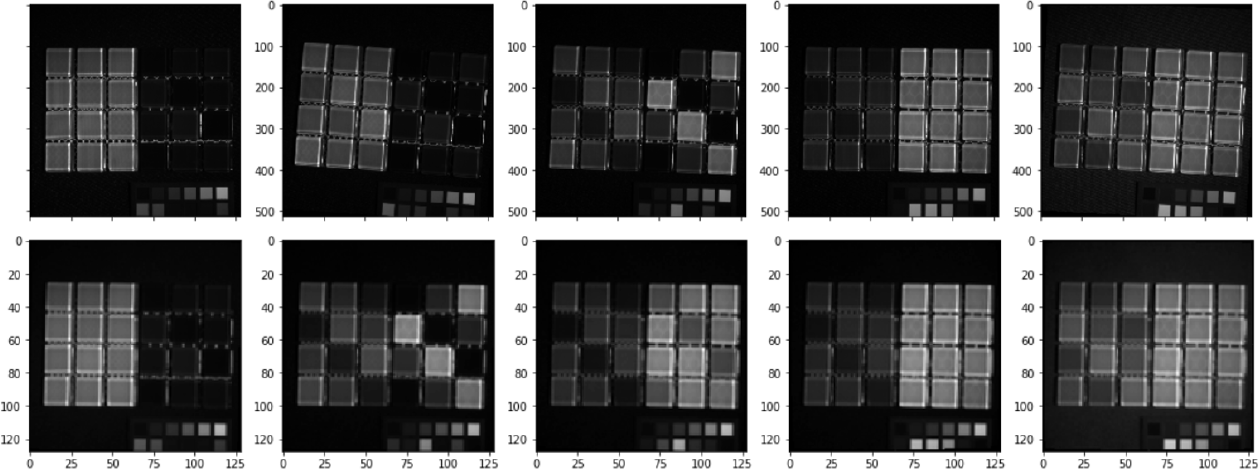
Top: Rotated test images. Bottom: Images registered using MM.

**Fig. 4.**
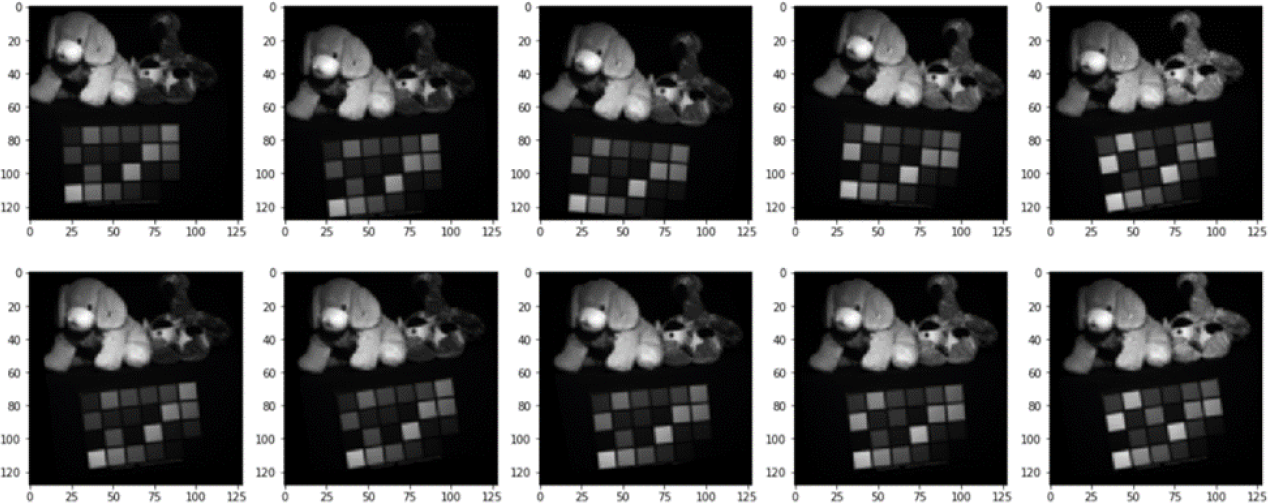
Top: Rotated and shifted test images. Bottom: Images registered using MM.

We also test a single maximum projection smFISH image; rather than resizing, we crop images and perform registration on areas with high densities of fluorescent markers. To increase the speed of PSO convergence, we use a dictionary to store the result of every computation so that the same calculations need not be repeated, and we introduce a spread factor to the PSO to improve convergence speed [34].

### B. Finite rate of innovation sampling

If the weighted spikes corresponding to innovative features are correctly located across each image, the sampled spike images will be transformed versions of the reference image, with transformations **T**_*δ*_ that are exactly the original transformations **T**. If a spike *δ*_*i*_ has moved an *L* − 2 distance less than *D* from the reference spike *δ*_1_ for all images *i* = 2,…, *n*, and if any spike corresponding to a feature is significantly greater than distance *D* from all other spikes in the image (that is, spikes are far apart and shifts and rotations were small), then we can use a nearest neighboring clustering algorithm to group spikes corresponding to the same feature across images.

We first demonstrate that we can register a collection of randomly shifted spikes. We randomly generate a 1000 × 1000 pixel image of 500 maximum intensity pixels (spikes) on a black background. We translate the image and use a nearest neighbor search to find where a spike has shifted. Both our algorithm and phase correlation register the images correctly with no error. We then add noise by randomly removing 50 spikes and randomly introducing 50 spikes. We then randomly shift individual spikes by 1 pixel to any of its eight adjacent locations. We find that both our algorithm and phase correlation produce an error of 1 pixel in any direction for about half of the images.

We test registration of multispectral images by randomly applying a random horizontal and vertical shift drawn from [−25, 25], and we sample using Rosten and Drummond’s fast corner detection [35]. We represent corner locations as spikes (see Fig. 5) and perform a nearest neighbor search using a search neighborhood of 30. FRI sampling followed by clustering again registers the images correctly around 40% of the time, and is off by 1 pixel in any direction 60% of the time. On the other hand, cross-correlation gives poor results when the images look drastically different in the various spectra. Fig. 7 shows a typical example when MM and our FRI method correctly register images, but phase correlation does not.

**Fig. 5.**
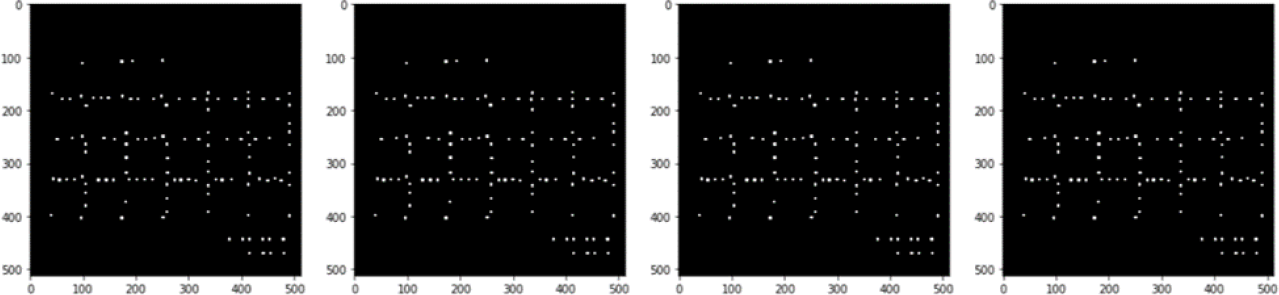
spikes found by corner detection.

Next we find a sampling procedure that works well with smFISH images, taking fluorescent markers as features. We begin again by performing a maximum intensity projection across all three color channels to obtain one image per imaging round. We locate the fluorescent markers using a White Top Hat filter, which extracts small points that are brighter than their surrounding and threshold to remove any low-intensity points generated by noise.

Since each fluorescent marker is several pixels large, we must find a single pixel location that best approximates each fluorescent marker. We use density-based spatial clustering of applications with noise (DBSCAN) since the number of clusters does not need to be specified and since it does not cluster outliers [36]. DBSCAN clusters pixels corresponding to a single fluorescent marker (see Fig. 6). Once clusters are identified, we find the coordinates of their centroids.

**Fig. 6.**
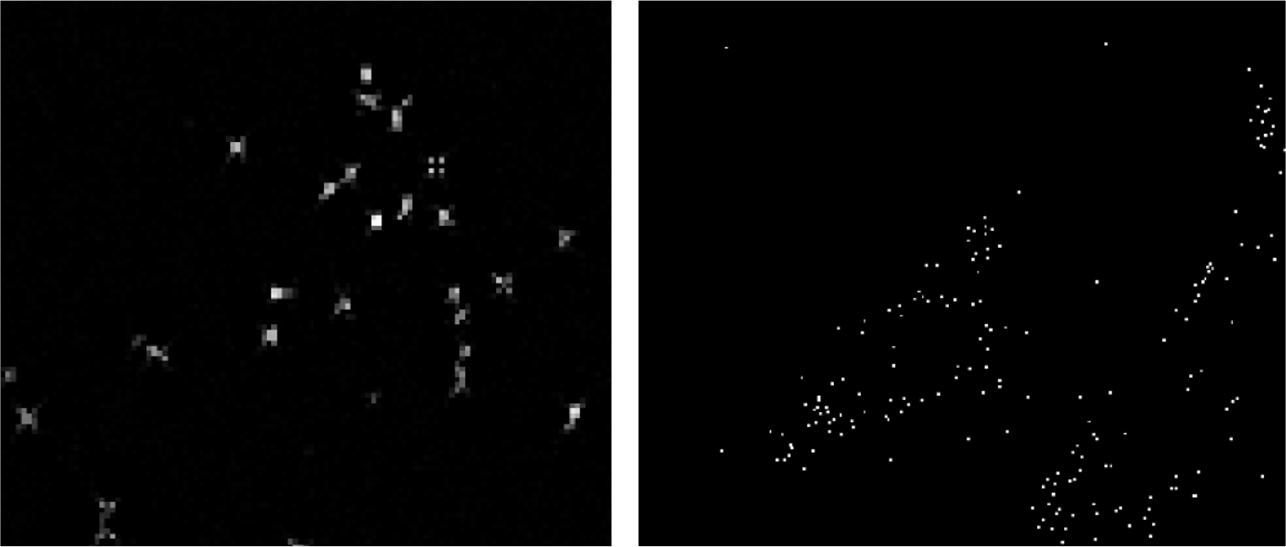
Left: Groups of pixels corresponding to fluorescent markers (enlarged for visibility). Right: spike deltas corresponding to centroids of clusters

**Fig. 7.**
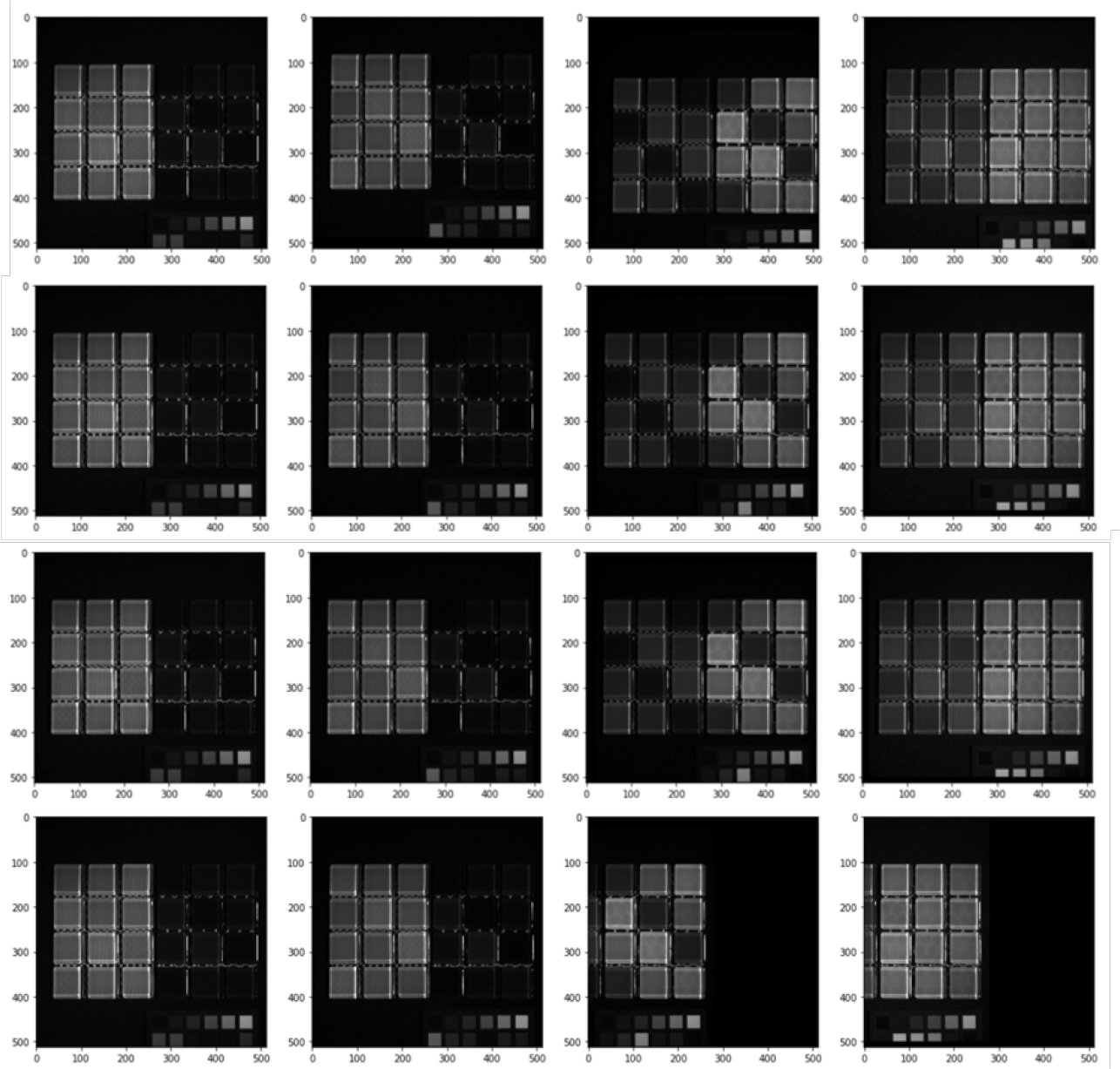
From top to bottom: Manually shifted test images, registration using MM (correctly registered), registration using FRI sampling (correctly registered), registration using phase correlation (incorrectly registered).

We obtain a list of reference spikes from the reference image and we align the spikes in each misaligned image to the reference spikes using nearest neighbor clustering across images. We find that results are more accurate when the search range for each point was small, so we use a successive refinement of the search radius. We perform the search first with a larger search radius of 50 and then with a smaller search radius of 25.

### C. Experimental Results

Registration results for smFISH images are shown in Figs. 8 and 9. Since there is no ground truth, we closely inspect image regions to visually confirm correct registration (see Fig. 10). Comparing our results to results using phase correlation indicate improved performance. Registration of four 980 × 1330 images takes under a second with both FRI and phase correlation and 15 minutes with MM on the same machine.

**Fig. 8.**
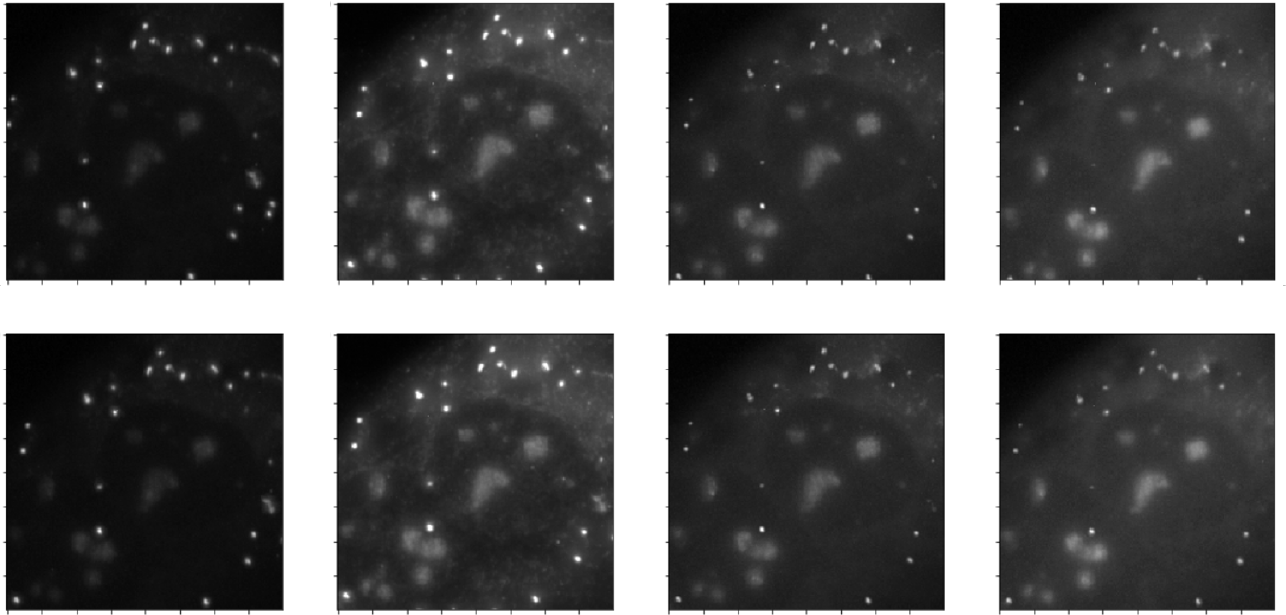
Registration results. Top: Before registration. Bottom: After registration with FRI

**Fig. 9.**
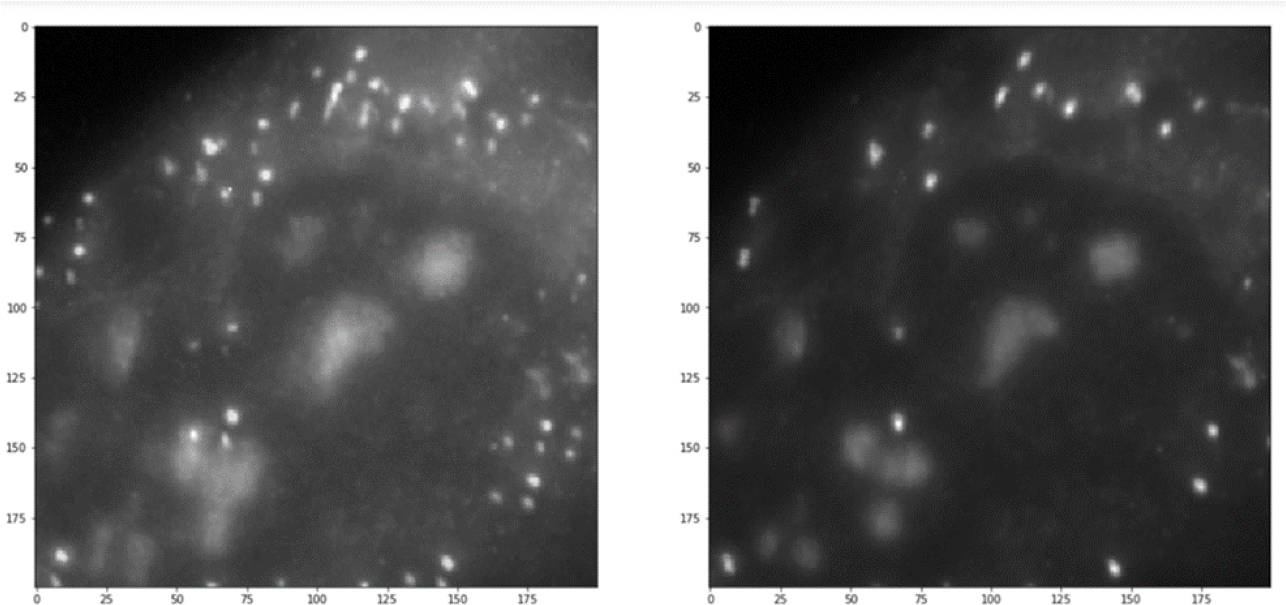
Left: Overlay of image rounds before registration. Right: Overlay of image rounds after registration with FRI.

**Fig. 10.**
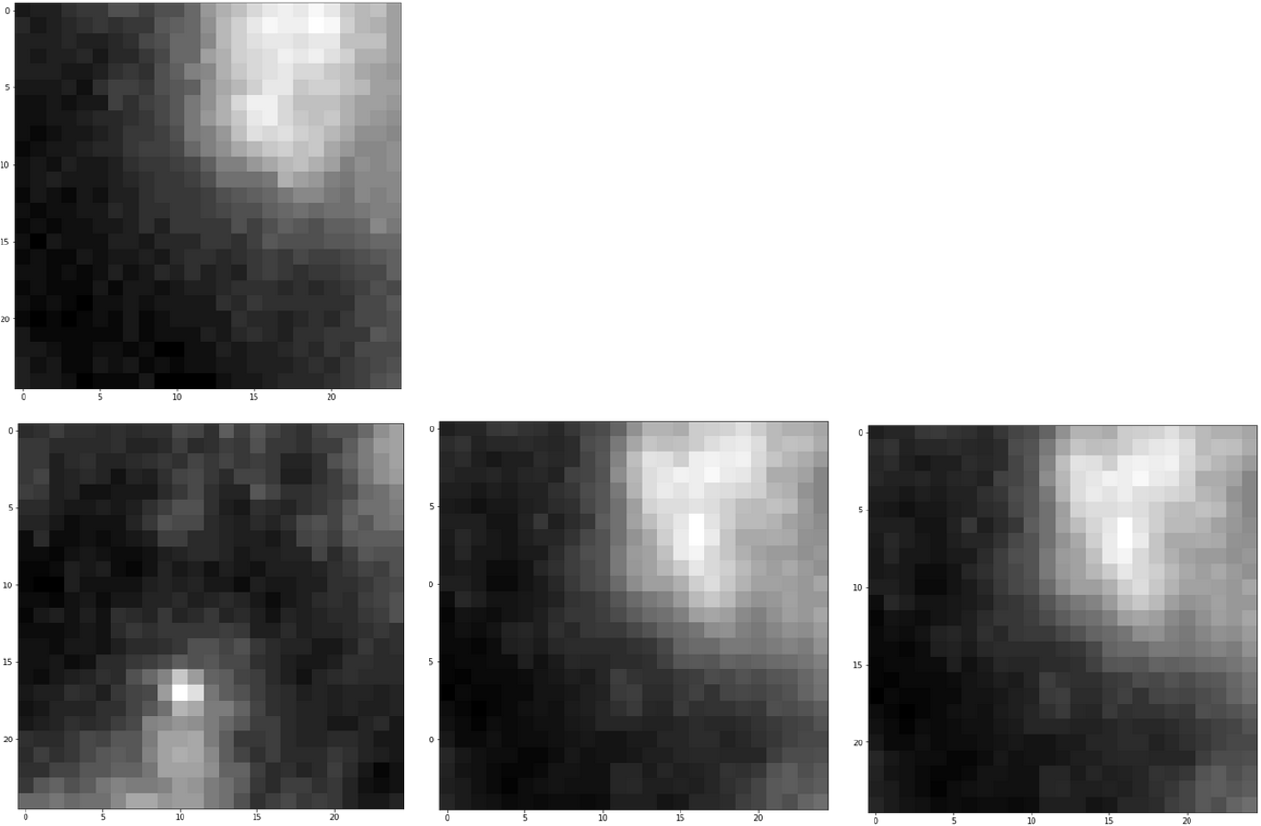
Close of up pairwise registration results. Top: Reference image. Bottom left: Before registration. Bottom center: After registration with MM. Bottom right: After registration with FRI.

## V. Discussion

Not only does finite rate of innovation sampling improve computational efficiency of groupwise image registration, as we have shown, but it is asymptotically optimal for image registration if it is used with multiinformation.

### Theorem 1.

*If images X*_1_,…, *X*_*n*_ *are FRI signals*, *maximizing multiinformation of the images sampled at FRI is asymptotically optimal for image registration*.

*Proof:* The maximum likelihood estimator of image transformations when transformations are independent is given by:

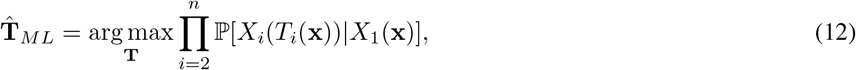

which is Bayes optimal. The MM estimate for image registration

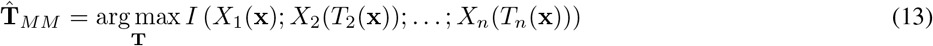

is exponentially consistent with 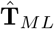, as we showed in [11, Theorem 7]. Let 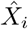 be *X*_*i*_ sampled at or above the rate of innovation for *i* = 1,…, *n*. Then there is no loss of information from *X*_*i*_(*T*_*i*_(x)) to 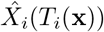 [12], and

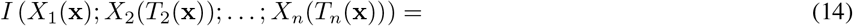

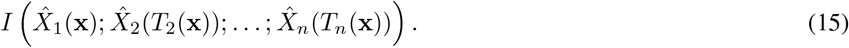

Thus 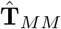 of the FRI-sampled images is equivalent to 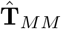 of the original images, which is asymptotically optimal.

Although MM is theoretically optimal, multiinformation is costly to compute even for FRI-sampled signals, so in practice we have used nearest neighbor clustering of sampled images. We have demonstrated this heuristic approximation is effective. Going forward, we aim to examine efficacy and computational efficiency of our technique for image-based transcriptomics at the massive scale of the Human Cell Atlas.

## Acknowledgment

We thank Deep Ganguli (Chan-Zuckerburg Initiative) and Ravi Kiran Raman for their help.

